# Multitrophic higher-order interactions modulate species persistence

**DOI:** 10.1101/2021.11.18.469079

**Authors:** Lisa Buche, Ignasi Bartomeus, Oscar Godoy

**Affiliations:** Departamento de Biología, Instituto Universitario de Investigación Marina (INMAR), Universidad of Cádiz, 11510-Puerto Real, Spain; Estación Biológica de Doñana (EBD-CSIC), Sevilla, Spain

**Keywords:** coexistence, multitrophic interactions, per capita effects, plant-pollinator network

## Abstract

There is growing recognition that interactions between species pairs are modified in a multispecies context by the density of a third species. However, how these higher-order interactions (HOIs) affect species persistence remains poorly understood. To explore the effect of HOIs steaming from multiple trophic layers on plant persistence, we experimentally built a mutualistic system containing three plants and three pollinators species with two contrasting network structures. For both structures, we first estimated the statistically supported HOIs on plant species, in addition to the pairwise interactions among plants and plant-pollinators. Following a structuralist approach, we then assessed the effects of the supported HOIs on the persistence probability of each of the three competing plant species and their combinations. HOIs produced substantial effects on the strength and sign of per capita interactions between plant species to such an extent that predictions of species persistence differ from a non-HOIs scenario. Changes in network structure due to removing a plant-pollinator link further modulated the species persistence probabilities by reorganizing per capita interaction strengths of both pairwise interactions and HOIs. Our study provides empirical evidence of the joint importance of HOIs and network structure for determining the probability of species to persist within diverse communities.

## 1 Introduction

Much of our understanding of the mechanisms that maintain species diversity comes from the study of pairwise direct interactions (Chesson, 2000; Adler et al., 2007) or the structure of pairwise interaction chains (Levine et al., 2017). These are chains such as rock-paper-scissors dynamics in which species are embedded in ecological networks but the nature of interactions is still between pairs of species (May and Leonard, 1975; Allesina and Levine, 2011; Godoy, Stouffer, et al., 2017). However, when addressing a multispecies context, the nature of interactions (i.e. strength and sign) between pairs of species can also be modified by the presence of a third species in the community (Billick and Case, 1994). For instance, tall plant species can cast shadows over other smaller plant species inducing plastic morphological or physiological responses (e.g. changes in foliage surface or light use efficiency) (Perez-Ramos et al., 2019; Kleinhesselink et al., 2019), which in turn can change the strength and even the sign of their interactions with other species. Similarly, the mere presence of a top-predator can change the behavior of their prey, and hence, the competition for shared resources among prey species (Wissinger and Jill, 1993; Beckerman et al., 1997). These interaction modifications are initiated by the presence of a third species (the initiator) which changes the per capita effect of a competing species (the transmitter) on a focal species (the receiver) (fig.1.a-b) (Li, Mayfield, et al., 2021).

**Figure 1:**
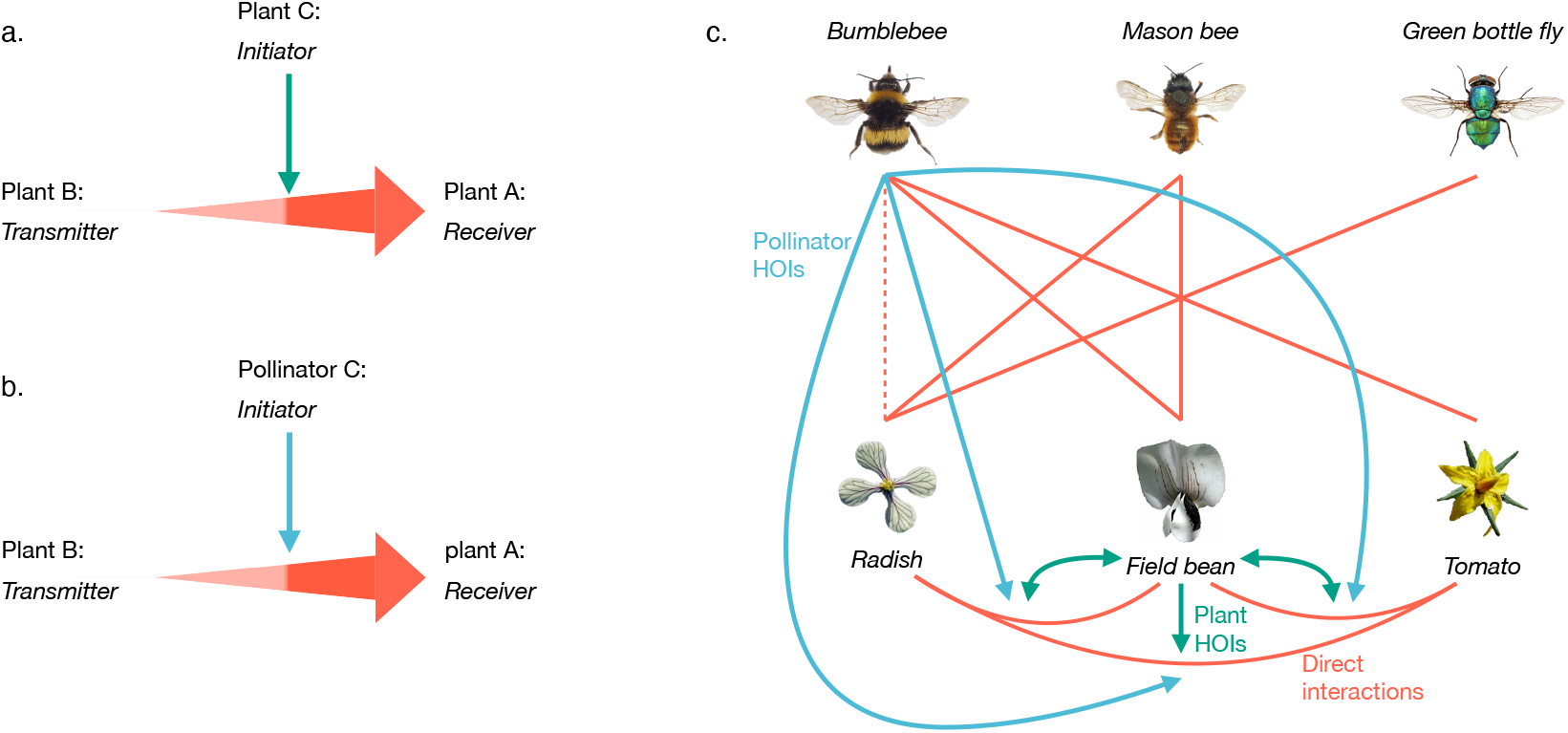
Interaction modifications, or “Higher-order interactions” (HOIs), are initiated by the presence of a third species (the initiator) which changes the per capita effect of a competing species (the transmitter) on a focal species (the receiver), ultimately changing the nature of the per capita interactions (i.e. strength and sign). **(a)** A plant HOIs implicates that a plant (the initiator) changes the phenotype of a plant B (the transmitter) and hence, the competition for shared resources, that is the nature of the per capita interaction among plant B and plant A. Whereas, in **(b)**, the initiator is a pollinator, hence we speak of pollinator HOIs. In both cases, we observe an HOI rather than an interaction chain, as we do not observe a direct change in the density, but a change in the strength and sign of the interaction itself **(c)** Plant-pollinator network under a complete and modified nested structure. The red lines show the direct interactions, either between two plants, i.e. plant pairwise interaction (*α_ij_*) or between a pollinator and a plant, i.e. mutualistic interaction (*γ_il_*). The dotted red line shows the pollinator-plant interaction removed to test the influence of the network structure. The green and blue lines designate respectively plant (*β_ijk_*) and pollinator HOIs (*β_ijl_*). The HOIs considered here are interaction modifications that change the nature of plants per capita interaction. Species *i* and *j* are always plant species.

Ecologists have long hypothesized that these interaction modifications, or “Higher-order interactions” (HOIs), are a common phenomena in both competitive (Case and Bender, 1981; Worthen and Moore, 1991; Billick and Case, 1994; Wissinger and Jill, 1993; Mayfield and Stouffer, 2017) and food web contexts (Morin et al., 1988; Wissinger and Jill, 1993; Paterson et al., 2015). But it has not been until recent years that researchers have systematically explored how these HOIs-mediated changes in per capita interactions modulate the dynamics and the functioning of ecological communities (Bairey et al., 2016; Grilli et al., 2017; Letten and Stouffer, 2019; Kleinhesselink et al., 2019; Singh and Baruah, 2019; Li, Bearup, et al., 2020). To date, most efforts are theoretical, partly due to the difficulty in gathering rigorous datasets to quantify HOIs. The two empirical studies of this sort have shown that including HOIs in competitive plant networks improves the predictive power of individual fitness models (Mayfield and Stouffer, 2017; Li, Mayfield, et al., 2021). Yet how these results further influence the stability of ecological communities remains unknown because empirical estimations of HOIs’ effects on per capita interactions have never been linked to the probability of species persistence (Saavedra, Medeiros, et al., 2020). Obtaining this knowledge is essential if we want to better inform future theoretical work of what is a plausible range of ecological possibilities as well as provide detailed expectations for further empirical research.

An important step to evaluate the effect of HOIs on the persistence of species embedded in multitrophic communities requires understanding first when HOIs emerge, and second how these HOIs are modulated by the structure of species interactions. As mentioned before, HOIs can emerge from multispecies interactions within and between trophic levels (i.e. the initiator and/or the transmitter can be from the same or a different trophic level than the receiver) (Wissinger and Jill, 1993; Bairey et al., 2016), yet their relative magnitude has never been compared before. Moreover, there is increasing evidence that species reorganize their interactions strength (CaraDonna et al., 2017; Bartley et al., 2019) depending on their abundances and a series of biological constraints such as species’ traits, evolutionary histories, or environmental factors (Cattin et al., 2004; Montoya et al., 2006; Morales-Castilla et al., 2015) but to what extent this reorganization also affects HOIs remains also unknown. We, therefore, need to establish these empirical linkages to better understand the net effect of multiple sources modifying the per capita interaction strengths that ultimately determine the stability of ecological communities (Neutel et al., 2007; Godoy, Bartomeus, et al., 2018).

Here, we undertook a controlled experiment of a plant-pollinator system using two different network structures (see Fig.1.c) as a first step towards quantifying HOIs in a multitrophic context, and then we revealed their effects on the probability of species to persist. Specifically, we first characterized the system by assessing competitive interactions among plant species and the mutualistic interaction of pollinators with plants. On top of this network structure, we used a framework based on individual fitness models (Mayfield and Stouffer, 2017) to quantify statistically relevant HOIs within one trophic level (Fig.1.a), here named “plant HOIs” (i.e. the initiator, the transmitter and the receiver are plant species) and between two trophic levels (Fig.1.b), here named “pollinator HOIs” (i.e. the receiver and the transmitter are plant species but the initiator is a pollinator species). According to previous work, we can expect a pollinator HOI when a pollinator triggers a phenotypical plant response such as the modification of flower duration or the composition of floral volatile organic compounds (Vivaldo et al., 2017; Burkle and Runyon, 2019). Finally, we use recent advances following a structuralist probabilistic approach to predict the effect of HOIs on the persistence probability of each plant species individually, and in all pair and triplet combinations (Saavedra, Medeiros, et al., 2020). With such approach coupling experiments with theory and associated mathematical tools, we were able to answer the following three questions: (i) Does the trophic role of the third species (plant versus pollinator) explain the magnitude of the effect of HOIs on changing per capita interactions? (ii) How do HOIs impact the probability of each plant species to persist as well as their combination? (iii) What is the role of network structure in reorganizing the effect of HOIs on the plants’ direct per capita interaction and ultimately plant persistence probabilities?

Overall, we found that including HOIs drastically changed our expectation of which species are likely to persist compared to the non-HOIs scenario. Plant and pollinator HOIs showed contrasting magnitudes on modifying per capita interactions. Specifically, the positive effect of pollinator HOIs on community stability was masked by the overwhelming influence of plant HOIs. Our experimental design also allowed us to discover that network structure further modulated the effect of HOIs on plant per capita interactions and species persistence probabilities. However, both network structures show different predictions when HOIs were not considered. Most of these results could only be obtained by combining theory with detailed experiments, which highlights the necessity to obtain empirical data to better understand how species can persist within diverse ecological systems.

## 2 Material and methods

### 2.1 Experimental set up

We designed a multi-species experiment to evaluate the effect of plant and pollinator HOIs (i.e. how the presence of a third species, the initiator, changes the per capita effect of a competing species, the transmitter, on a focal species, the receiver (Li, Mayfield, et al., 2021)) on changing the nature (strength and sign) of per capita interactions among plant species pairs. To do so, we used ecological knowledge to create a plant-pollinator mesocosm composed of three plant species (*Raphanus sativus* (Brassicaceae): Radish, *Solanum Lycopersicon* (Solanaceae): Tomato, *Vicia faba* (Fabaceae): Field bean), and three pollinator species (*Bombus terrestris* (Apidae): Bumblebee, *Osmia bicornis* (Megachilidae): mason Bee, *Lucilia sericata* (Calliphoridae): green bottle Fly). All plant species are suitable for our experiment because they depend on pollinators to maximize seed sets. These species were also chosen to assemble a fully nested system in which flower and pollinator morphology determine species interactions (Fig.1). Therefore, we obtained a gradient from a generalist pollinator species, the Bumblebee, which can pollinate the three plant species including Tomato by buzz pollination, to a specialist species, the green bottle Fly, which can only visit the open flowers of Radish. The mason Bee can visit Radish flowers and it is also big enough to open the corolla of Field bean. Likewise, Radish was the generalist plant species in our system, which was visited by all pollinators, and Tomato was the specialist species, in which flowers were visited only by Bumblebee. Field bean individuals were visited by both the Bee and the Bumblebee species (Appendix, section 5.1, Fig.S2). To create a variation in this fully nested structure, we experimentally removed the link between Bumblebee and Radish by physically preventing their mutualistic interaction using a large mesh size (9 mm) that excludes Bumblebee but allowed the visitation of Radish flowers by the other two smaller pollinators (Appendix, section 5.1, Fig.S1).

To empirically evaluate the effect of plant HOIs (i.e. the initiator, the transmitter, and the receiver are plant species) on changing the nature of per capita interaction of one plant competitor on another, we need independent variation in the three species’ densities. The per capita interaction measures how much fitness (measured in our case as viable seed production) of a given species decreases (competition) or increases (facilitation) on average as a function of the density of the second species. Moreover, to evaluate how sensitive are each of the three plant species to plant HOIs, we additionally require variation in the relative density of the third species, the initiator. To account for these two sources of variation, we assembled communities of the three plant species in pots of 150 lts following an explicit spatial arrangement (Appendix, section 5.1, Fig.S1), in which individuals of each of the three species were specifically located to be surrounded by an increasing number of neighbors from 1 to 4 within a radius of 7.5 cm (a distance at which annual species are known to interact) (Godoy and Levine, 2014; Mayfield and Stouffer, 2017). Each pot contained a total of 23 individuals of the three plant species in different proportions. We also grew individuals in pots with no neighbors (9 individuals per plot spaced at least 15 cm) to empirically estimate the intrinsic growth rate of each species’ population in the absence of plant-plant interactions in these experimental conditions.

To evaluate the effect of pollinator mutualistic interactions on plant fitness, as well as pollinator HOIs (i.e. the receiver and the transmitter are plant species but the initiator is a pollinator species) on changing per capita interactions between plant species, we assembled 17 sealed cages of 3 *m*^3^. These cages were equally divided into different pollinator treatments in which we modified the density of each pollinator alone or in a combination of two pollinators together (including the case of no pollinator presence, Appendix, section 5.1, Fig.S1 for the number of individuals per cage). All pollinators nested within the cages (see Bartomeus et al. (2021) for details). For Bumblebees, we manually removed newly emerged workers to keep the number of individuals constant. Each cage contained either five or 10 plant pots (i.e. 8 pots for competition and 2 pots for no competition), which totaled 170 pots.

We ran this experiment over the growing seasons 2016-2017 at IRNAS-CSIC facilities (Finca La Hampa, http://www.irnas.csic.es/en/nca-experimental/), located in southern Spain (37°17’01.4”N 6°03’56.5”W). Five viable seeds per species were sown at each specific location within a pot to ensure individual presence. As soon as they germinated during winter, they were thinned to a single individual per pot location. In total, we measured the seed set on 907 individuals. On average, Radish plants produced 41.2 ± 42.5 seeds (mean ± standard deviation), Field bean plants produced 4.9 ± 3.1 seeds, and Tomato plants produced 74.5 ± 64.8 seeds.

### 2.2 Quantification of HOIs

Thanks to the experimental approach described above, we quantified the magnitude of plant and pollinator HOIs using the framework developed by Mayfield and Stouffer (2017). This framework is based on a model that describes, using a negative exponential function, the average decay of individual fitness as a function of density-dependent processes arising from pairwise interactions and HOIs. The general description of this model is the following:

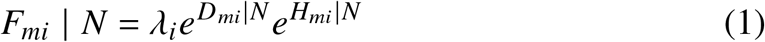

with *F_mi_* the specific fitness of an individual *m* of the species i for a specific set *N* of neighbors, *λ_i_* the intrinsic fitness of species *i* in the absence of competitors, and the exponential terms correspond respectively to the ensemble of pairwise interactions *D_mi_* and HOIs *H_mi_* for a specific set *N* of neighbors.

Here we extended this general form to a multi-trophic context in which we included the mutualistic effect of pollinators on plant species, and we further specified whether HOIs acting on plants stem from a plant or a pollinator species. Accordingly, the cumulative effect of the pairwise interactions of the set of neighbor species *N* follows the standard linear density dependant form:

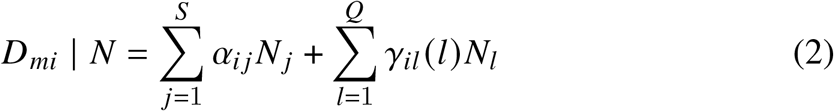

where, *N_j_* is the abundance of species j, *α_ij_* is the direct effect of species j on the focal species i, and the sum is across all the neighbor plant species *S* specific to species *i*, *γ_il_*(*i*) is the functional response of the pollinator species *l* on the focal species *i*, and the sum is across the pollinator species *Q* specific to species *i*. *α* can be positive or negative, capturing respectively, facilitation or competition. *γ*(*l*) can be of linear function (i.e. type I) or a noninflected functions (i.e. type II) (see section5.5). *γ*(*l*) represents the mutualistic interaction, which positively affects the intrinsic fitness of the focal species *i*.

The ensemble of HOIs acting on species *i* is quantified according to the following equation:

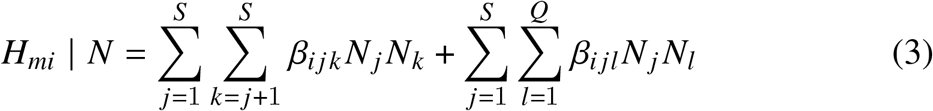

*β_ijk_* is the effect of a plant population, species *k*, on the per capita interaction between two plants, species *i* and species *j* (here *β_ijk_* is called “plant HOIs”), with *j* ≠ *k*, *β_ijl_* is the effect of a pollinator population, species *l*, on the per capita interaction between two plants, species i and species *j* (here *β_ijl_* is called “pollinator HOIs”). For both type of HOIs, the effect is cumulative across the plant *S* and/or pollinator *Q* neighbours specific to the focal species *i*. *β_ijk_* and *β_ijl_* can change the strength and sign of the per capita interaction of a competitor on the focal species *i*. For each plant species, a maximum likelihood approach allowed the estimation of each coefficients.

Here, *N* was either the abundance of each of the plant species within the neighborhood or the number of pollinator visits observed for a specific period of time. To obtain this last piece of information, we measured the visitation rates of each pollinator to each plant in 3 minute periods twice during peak bloom for each cage (total = 576 min of observations). Additionally, pollen deposition per visit for each plant-pollinator combination was measured to ensure all pollinators are effective following Winfree et al. (2008). Mean pollen deposition per single visit was not different across species (*p* > 0.1).

### 2.3 Selection of pollinator functional responses and HOIs

To evaluate the importance of HOIs for the fitness of each plant species, we constructed three different types of models: (i) “Direct interactions” or “non-HOIs”, which means that *H_mi_* were not included, (ii) “Selected HOIs”, *H_mi_*, which means that only statistically supported HOIs (*βs*) were included, and finally, (iii) “All HOIs”, *H_mi_* which included all HOIs (*βs*) regardless of their statistical support. The three model categories always included the direct pairwise interactions between plants (*α*) and between plants and pollinators (*γ*). Regarding plant-pollinator interactions (*γ*), prior work has shown that the functional responses of a plant reproductive success to an increasing visitation rate of a pollinator (*l*) can take a linear shape, (type I: **γ**(*l*) ~ **γ*_i_l*) or noninflected shape (type II: *γ*(*l*) ~ 1/(1 + *γ_il_*) based on the cost-benefit ratio induced by the pollinator (Morris et al., 2010). Thus, before accounting for HOIs during our modeling procedure, we first selected the most parsimonious model accounting for pairwise interactions based on the Akaike information criterion (AIC, best-supported models are those with lower AIC), which includes either the type I or the type II response for the effect of each pollinator species on the fitness of each focal plant species (via the *dredge(MuMIn)* function). We believe this procedure ensures that we do not later attribute explained variance to pollinator HOIs (*β_ijl_*) that can be instead explained by a particular pairwise plant-pollinator functional response.

Finally, to further include HOIs (both plant and pollinator HOIs) into the “All HOIs” models, we departed from the best-supported models of pairwise interactions to which we added the estimation of all potential HOIs. Then, we obtained “Selected HOIs” models by including a subset of HOIs leading to the smallest AIC. The inclusion of a subset of HOIs always reduced AIC for all focal species compared to including only direct pairwise interactions between plants (*α*) and between plants and pollinators (*γ*), that is the non-HOIs model (Appendix 5.5, Table.S2). When multiple HOIs models were equally supported by AIC (i.e. Delta-AIC < 2), we separated them by looking at the highest log-likelihood value following (Mayfield and Stouffer, 2017) (Appendix, section 5.5). On average, across focal plant species, the “Selected HOIs” models included a subset of 3.66 ± 0.33 of both plant and pollinator HOIs, that is from 3 to 4 HOIs (Appendix, section 5.5, Table.S2).

### 2.4 Evaluation of the relative importance of plant and pollinator HOIs for changing the structure of plant interactions

By using the individual fitness model (Eq.1) for the three plant species, we obtained a matrix of plant interactions. This matrix named matrix A (see below) varied according to the three categories of the model considered (non-HOIs, Selected HOIs, and All HOIs). For each model, the matrix containing the per capita interaction was computed based on the different elements included in the *D_mi_* and *H_mi_* (see Eq.3). To evaluate the relative importance of plant and pollinator HOIs for changing the structure of plant interactions, we differentiate between plant and pollinator HOIs in *H_mi_*. Theoretically, both plant and pollinator HOIs have equal potential to modify a plant pairwise interaction, which we evaluated by the addition of HOIs on the plants per capita interaction computed in *D_mi_* (see Fig.2).

**Figure 2:**
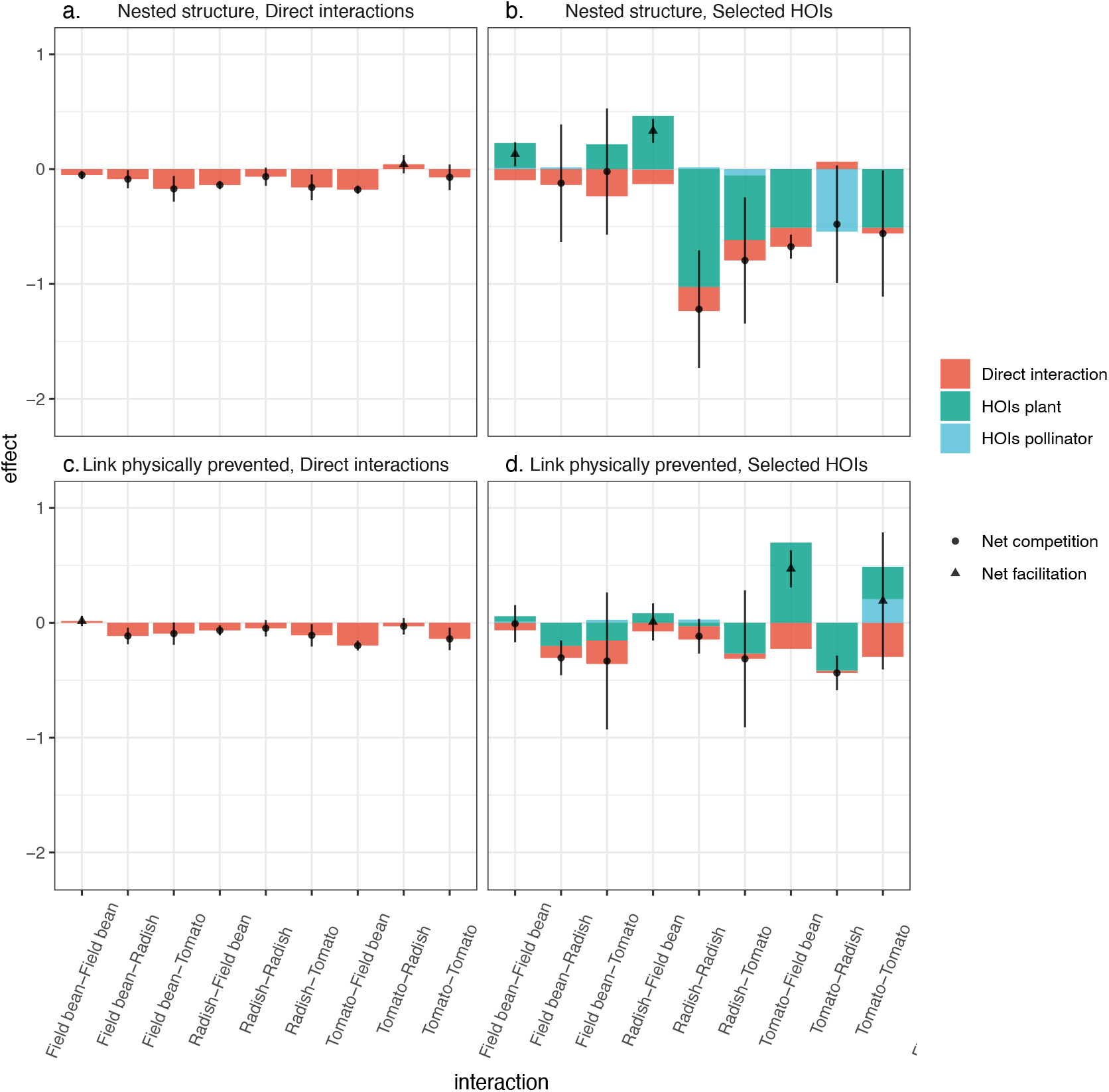
HOIs change the per capita strength and sign of the interactions between plant species pairs. These changes are modulated by the network’s structure. The nine direct plant interactions for the four scenarios are divided in terms of direct interaction(s) (red), plant HOIs (blue), and pollinator HOIs (green). The dots represent the sum of all interactions (i.e. the realized per capita interaction), that is the value of the resulting interaction strength and sign between the two corresponding plant species. The dot is circular if the value is negative (i.e. net competition) and triangular if the value is positive (i.e. net facilitation). Each dot is associated with its standard deviation.

### 2.5 Effect of HOIs on species persistence

We followed a structuralist probabilistic approach (Song, Rohr, et al., 2018; Saavedra, Medeiros, et al., 2020) to evaluate the ecological effect of the inclusion of HOIs on species persistence. This structuralist probabilistic approach computes the persistence probability of a species for the three categories of the model (Appendix, section 5.3). Hence, we can evaluate whether species persistence changes based on the inclusion (or not) of HOIs in the distribution of the interaction matrix. We advance that predictions of species persistence are rather similar between the “Selected HOIs” and “All HOIs” models (Appendix, section 5.5, Fig.S7). Therefore, we present in the next section only results describing the effect of HOIs on species persistence of the “Selected HOIs” models.

Briefly, the evaluation of species persistence following a structuralist probabilistic approach works as follows: for a given matrix of interactions among three plant species *i*, *j*, and *k* (matrix A), this approach predicts the range of potential directions of vectors of intrinsic growth rates (r-vectors (*r_i_, r_j_, r_k_*)) compatible with the positive abundances at equilibrium (i.e., *N*\* = *A*^−1^*r*) of every single species as well as their combinations in pairs and the triplet. (Appendix for more details of this analysis, section 5.3). Accordingly, two or three species are predicted to persist together (i.e. coexist) when they share identical directions of r-vectors that fulfill the condition of positive abundances at equilibrium. In other words, two or three species are predicted to coexist when the structure of species interactions can accommodate differences in their intrinsic growth rates. For example, species that coexist because they can accommodate a large range of differences in intrinsic growth rates are those in which intraspecific interactions are stronger than interspecific interactions (Barabás et al., 2016).

In prior work using a structuralist approach, the matrix A containing per capita plant interactions is considered as a fixed element (Saavedra, Medeiros, et al., 2020). However, in our case the matrix A varies and it is drawn from the specific distribution of the interaction matrix. The distribution is based on the estimated elements included in the *D_mi_* and *H_mi_* (see Eq.3). The per capita interaction of two plants and their corresponding HOIs determine a specific distribution corresponding to their net interaction. That is, each element of an interaction matrix was computed as a normal distribution with a mean equal to the direct interaction and a variance equal to the sum of the HOIs. For instance, for the “All HOIs” model, the net interaction between the species i and the species j is distributed over 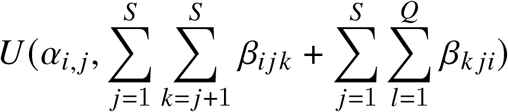. Similarly, the variance for the “Direct interactions” model is always zero, while the “HOIs selected” model only includes those HOIs that were statistically supported (see Appendix 5.5, Table.S2). Based on these distributions, we randomly chose 10 matrices A to evaluate each sample of the vector of intrinsic growth rates (see section below). In Fig.3, one location corresponds to one vector of intrinsic growth rates evaluate over 10 matrices A. Hence one location exhibit a dominant color, translating the species combination which has the highest probability.

**Figure 3:**
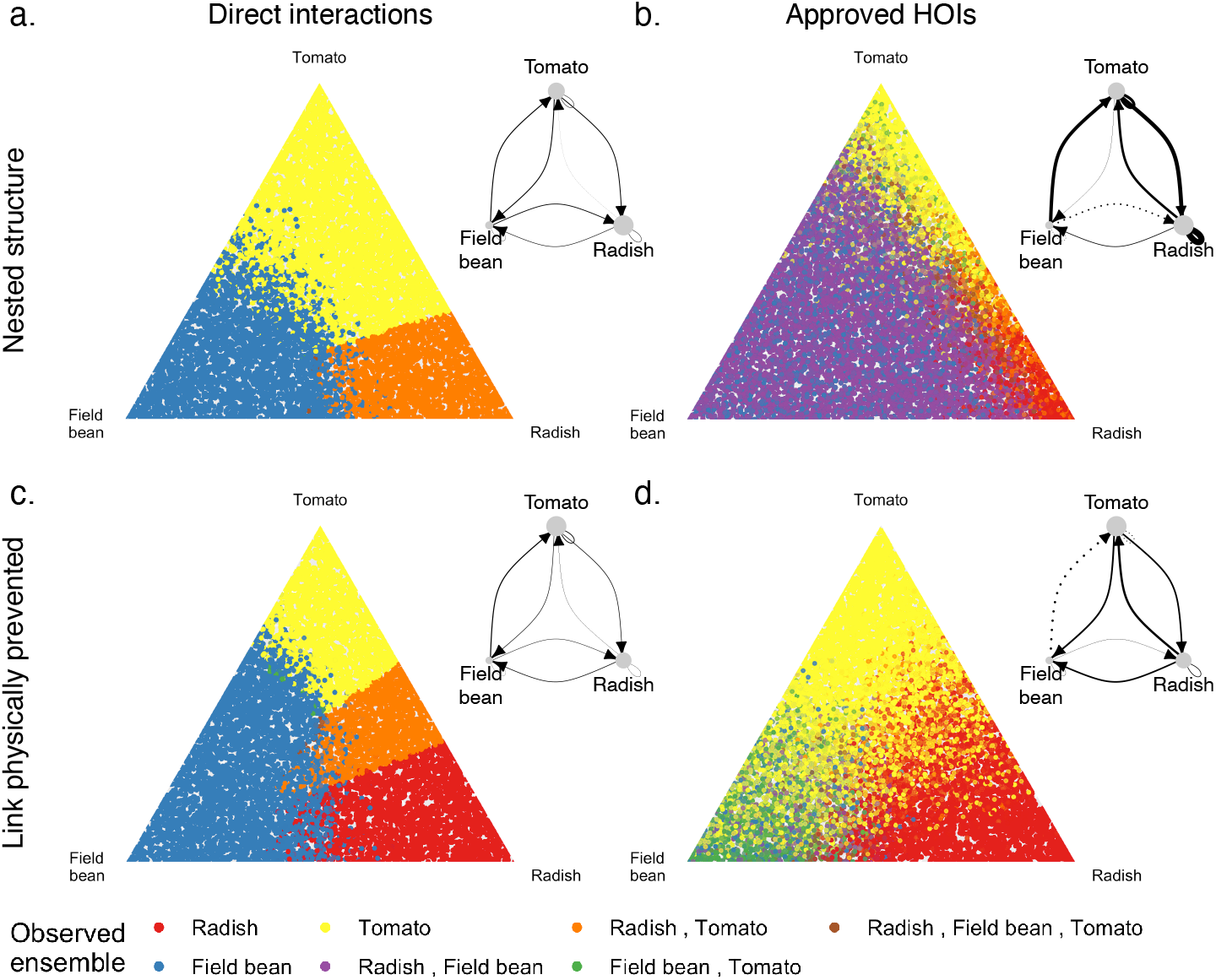
The inclusion of HOIs affects the probability of species persistence in both network structures. The percentage of persistence probability of each plant species individually, and in all pairs and triplet combinations is proportional to the fill of the triangle under the corresponding color. Each triangle is associated with its corresponding diagram of species interactions (solid lines = net competitive interaction, dotted lines = net facilitation and the width of the line is proportional to the absolute value of the strength of the interaction). **(a)** In a nested structure without HOIs, three combinations can be observed, of which two include only one plant species. **(b)** For a similar structure, the inclusion of HOIs increases the overall probability of pairs communities to persist, and especially the community of Radish and Field bean which is partly due to the net positive interaction between them. **(c)** When the link between the Radish and the bumblebee is prevented, with no HOIs included, the number of combinations increases of which the majority includes Radish. **(d)** For this structure, the inclusion of HOIs reduces the probability of pairwise community to persist due to the increase of individual persistence probability, particularly for Radish. The triplet combination is observed only in (b) which shows how the network structure determines the effect of HOIs on species persistence.

Moreover, to compare the cumulative effect of HOIs on changing the structure of the interaction matrices at the community level, we used a Procrustes analysis (also called a Procrustean superimposition approach) (D. Jackson, 1995; Peres-Neto and D. A. Jackson, 2001) (Appendix, section 5.2). Note that the HOIs including two or more pollinators are not considered here as they do not change the nature of the plant interactions.

## 3 Results

HOIs changed the per capita strength and sign of the interactions between plant species pairs (Fig.2). According to the “selected HOIs” models, for the nine species interactions contained in our matrix of 3 × 3 plants in the fully nested structure, HOIs increased competition (i.e. HOIs are negative) in five of the nine cases, and relaxed competition (i.e. HOIs are positive) in the other four cases. This reinforcement of positive interactions reversed two plant interactions from competition to facilitation (see interactions Radish - Field bean and Field bean - Field bean, Fig.2). Similarly, in a structure with the link Bumblebee-Radish physically prevented, HOIs changed the sign of four interactions between pairs of plants. Three of these changes were from negative to positive interaction signs (see interactions Radish - Field bean, Tomato - Field bean, and Tomato-Tomato, Fig.2).

The magnitude of changes implemented by plant HOIs was greater compared to the pollinator HOIs (Appendix, section 5.4, Fig.S5). Indeed, HOIs showed an effect stronger than the standard deviation of the direct interaction estimates in 77.8 to 100 % of cases (depending on the model and network’s structure) for plant HOIs and only 16.7 to 33.3 % for pollinator HOIs (Appendix, section 5.4, Table.S1). Yet, the relative percentage of positive pollinator HOIs was always greater than the relative percentage of positive plant HOIs (Appendix, section 5.4, Table.S1 and section 5.5 Table.S2 for the coefficients). Regarding individual species, we observed that Field bean was the less sensitive species to HOIs from other species, that is its interactions’ strength and sign were less impacted by the inclusion of HOIs. At the other extreme, Tomato was the species most sensitive to HOIs regardless of the trophic role of the third species. Note that Tomato is the species with slower development in terms of plant growth rates and Field bean the fastest.

As a consequence of changing the structure of species interactions (Appendix, section 5.2), HOIs increased the probability of more diverse communities persisting under the fully nested structure. While the feasibility domain accounting only for direct interactions between pairs predicted the persistence of single species (Field bean and Tomato separately) and one two-species combination (Radish and Tomato), the inclusion of HOIs predicted the persistence of two two-species combinations (Radish-Field bean, and Field bean-Tomato separately) and all species together. The inclusion of HOIs also enlarged the feasibility domain of the species pair Radish - Field bean, partly due to the net positive interaction between them.

Contrastingly, the empirical prevention of one mutualistic interaction (Radish and bumblebee) reorganized both the per capita interaction strengths and its modifications by HOIs to such extent that HOIs no longer promoted the persistence of two species (Radish and Field bean, Fig.3). Rather, they promoted the dominance of Tomato. In this case, the inclusion of HOIs decreased the individual persistence probability of Field bean (Appendix, section 5.3.1, FigS4). This decrease can be attributed to the translation of the unidirectional positive interaction between Field bean-Radish to Field bean-Tomato.

Overall, HOIs changed the persistence probability of each plant species individually, and of each pair and triplet combination under both network structures (Fig.3 and Appendix, section 5.3.1, Fig.S4). That is, the prediction of which combination of persisting species differed from the scenario of not considering HOIs (Fig.3). Specifically, Tomato, Field bean, and their combination were predicted to have higher chances to dominate the community in non-HOIs, than in a HOIs inclusive scenario (Fig.3). But Tomato was mostly substituted by Radish in the HOIs scenario. This final result suggests that the consideration of HOIs has profound implications for our ability to predict the composition of ecological communities.

## 4 Discussion

Higher-order interactions (HOIs) are common phenomena in nature that hold the potential to modify the strength, and even the sign, of per capita interactions in multispecies communities (Billick and Case, 1994; Mayfield and Stouffer, 2017; Levine et al., 2017). However, how these modifications further modulate predictions of species persistence has been underexplored in empirical settings. By combining individual fitness models and a structuralist probabilistic approach with detailed experiments, we provide three lines of evidence suggesting that we need to take into consideration trophic level and network structure information to better forecast the effect of HOIs on multispecies coexistence. First, we show that plant and pollinator HOIs simultaneously occur but with different magnitudes. Second, the network structure can reorganize HOIs strength, a result that can not be gauged using simulation approaches alone (Bartomeus et al., 2021). Third, HOIs arising from distinct network structures can result in contrasting effects on the probability of species and communities to persist. As a result, predictions of which combination of species are more likely to persist can change considerably depending on whether multitrophic HOIs are or are not considered together or separately.

We report that plant HOIs, in general, had a stronger effect than pollinator HOIs on changing per capita interactions between pairs of plant species (Appendix 5.4, Fig.S5). We believe these differences in magnitude are specific to our experiment and thus, it is early to suggest that they are representative of what can happen under more natural settings. Therefore, further studies including pollinators as well as other trophic interactions such as herbivory or soil biota organisms (e.g. mycorrhiza) are needed to confirm that HOIs coming from multitrophic interactions have a lesser impact than plant HOIs at modifying per capita plant interactions. Experimental settings might not be amenable to more complex systems, but with the analytical toolboxes already available, we can directly compare the relative importance of different trophic HOIs by accounting for the natural variation in multitrophic interactions and species relative abundances occurring along environmental gradients (e.g. (Lanuza et al., 2018; CaraDonna et al., 2017; Rumeu et al., 2019)). It is nevertheless clear from our experiment that such evaluations need to take into account the network of multitrophic interactions because alterations in their configuration can further modulate the effect of HOIs on species persistence (Fig.3). There were no previous expectations that the reorganization of interaction strengths after modifying our experimental network reduced the opportunities for species to coexist when we include HOIs (Fig.3). This result contrasts with a similar previous study showing that the modification of the network structure increases species persistence probabilities when HOIs are not considered (Bartomeus et al., 2021). These discrepancies between previous findings suggest that we need to reevaluate the effect of interaction strength and network structure on species persistence. Traditionally, it has been posited that highly nested plant-pollinator structures promote plant coexistence because they distribute interaction strength in such a way as to reduce competition between plant species (Bastolla et al., 2009; Verdú and Valiente-Banuet, 2008). However, mutualistic direct interactions and pollinator HOIs can also promote competitive dominance of some plant species over the others, which leads to species exclusion (e.g. when pollinators benefit the most competitive plants) (Bastolla et al., 2009; Godoy, Bartomeus, et al., 2018; Johnson, 2021). How different opportunities for species to coexist emerge under different network configurations due to the combined effects of direct, indirect, and high-order interactions is an avenue of research that deserves future attention.

Similar to prior work dealing also with annual plant species, we found that HOIs were species-specific (Mayfield and Stouffer, 2017). This variation between species could not be put in relation with any trait (Kleinhesselink et al., 2019), except with development speed. For instance, Field bean, the fastest-growing species in our experiment, experienced the smallest influence from all types of HOIs. At the other extreme, all HOIs were relatively stronger on Tomato, the slowest-growing species (Appendix 5.4, Fig.S5). Although our experimental setting differs from that of Mayfield and Stouffer (2017), both studies provide mounting evidence that HOIs do not have a net directional effect on favoring some species while harming others. Rather, they tend to increase the variance of interaction strength among species. An increase in random variance has been shown to reduce the stability of ecological communities, and change the composition of species predicted to persist for large species assemblages (Neutel et al., 2007; McCann et al., 1998). However, in our study, which is a low diverse system, the inclusion of the HOIs statistically supported increases rather than decreases the persistence probability of the complete plant community, that is, the three-species combination (Appendix 5.5, Fig.S7). Very similar results were obtained when considering all fitted HOIs (see Appendix 5.5, Fig.S6). Hence, our results provide empirical support to recent theoretical studies indicating that HOIs can stabilize the dynamics of interacting species (Grilli et al., 2017; Bairey et al., 2016). The transition to studying the effect of HOIs on species persistence from low to highly diverse systems can be challenging for two main reasons. First, the number of parameters to estimate scales exponentially with the number of species. This means that we take the risk of over-fitting models. Second, it is not straightforward to identify which subset of HOIs impact the dynamics of interacting species. To overcome these two limitations at once, recent statistical approaches are combining HOIs with information that can group individual species according to general categorizations (e.g. common vs rare species, grass/herb, slow/fast-growing) which ultimately, can reduce model dimensionality (Martyn et al., 2021). These approaches hold the ability to build stronger statistical models that allow the inclusion of HOIs for more species while ensuring ecological realism and model tractability.

An early approximation to the study of HOIs was to document how specific species can modify interactions between competitors (Wissinger and Jill, 1993). This approach was recently extended to include a multispecies context (Grilli et al., 2017; Mayfield and Stouffer, 2017). Here, we further enlarge the domain of HOIs establishing connections from populations to multitrophic networks thanks to a controlled experiment. However, we believe this is just the first step, and the challenge now remains to evaluate these empirical connections across spatial and temporal dimensions under natural conditions in which species interact under different network structures. Because HOIs permeate across levels of ecological organization, our results suggest that we need to explicitly account for them if we want to better understand the importance of species interactions for community dynamics.

## 5 Acknowledgement

We thank Prof. Margaret Mayfield for comments on an earlier version of the manuscript. We thank Semillas Fito for kindly providing the seeds for the experiment and to “Estación Experimental La Hampa” for access to the experimental facilities. Curro Molina and Mary Berry for helping with the experimental setup and data collection. This manuscript is funded by the LINCX project (CGL2014-61590-EXP). OG acknowledges financial support provided by the Spanish Ministry of Economy and Competitiveness (MINECO) and by the European Social Fund through the Ramón y Cajal Program (RYC-2017-23666).

## Appendix

### 5.1 Experimental design

**Figure S1:**
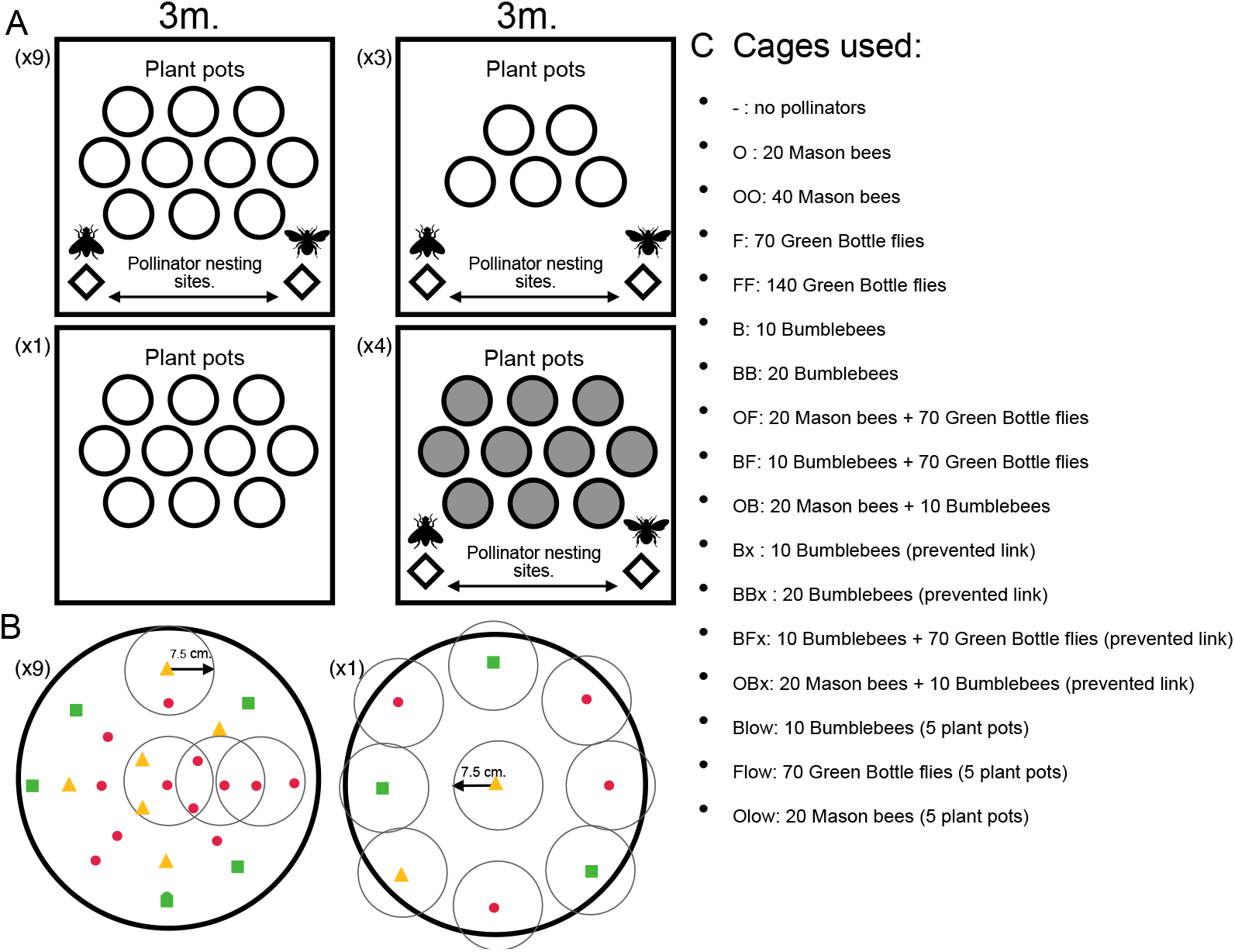
A: The four types of experimental cages included 4 to 10 different plant-plant competition pots, and zero or two pollinator nesting sites. The number of cages of each type is indicated respectively at the top left. B: Examples of the configurations of the three plant species (indicated by different colors and shapes) in the plant-plant competition pots. Each plant individual is surrounded by 4 to 1 plant neighbors (within a radius of 7.5 cm), except for two pots where plant individuals grew in no competition (no neighbors present in > 15 cm). Within a cage, the 10 pots had all three plants growing in different configurations that create all combinations of inter-and intra-specific competition at all densities. C: The 17 cages had different pollinator treatment levels that allow us to calculate the plants’ respective intrinsic growth rate, plant direct pairwise interactions, the plant HOIs and the pollinator HOIs, for both network structures. The number of pollinator individuals is selected to use the same pollinator biomass for each species (i.e. bumblebees are 7 times bigger and active than flies). For bumblebees, we manually removed newly emerged workers to keep the number of individuals constant.

**Figure S2:**
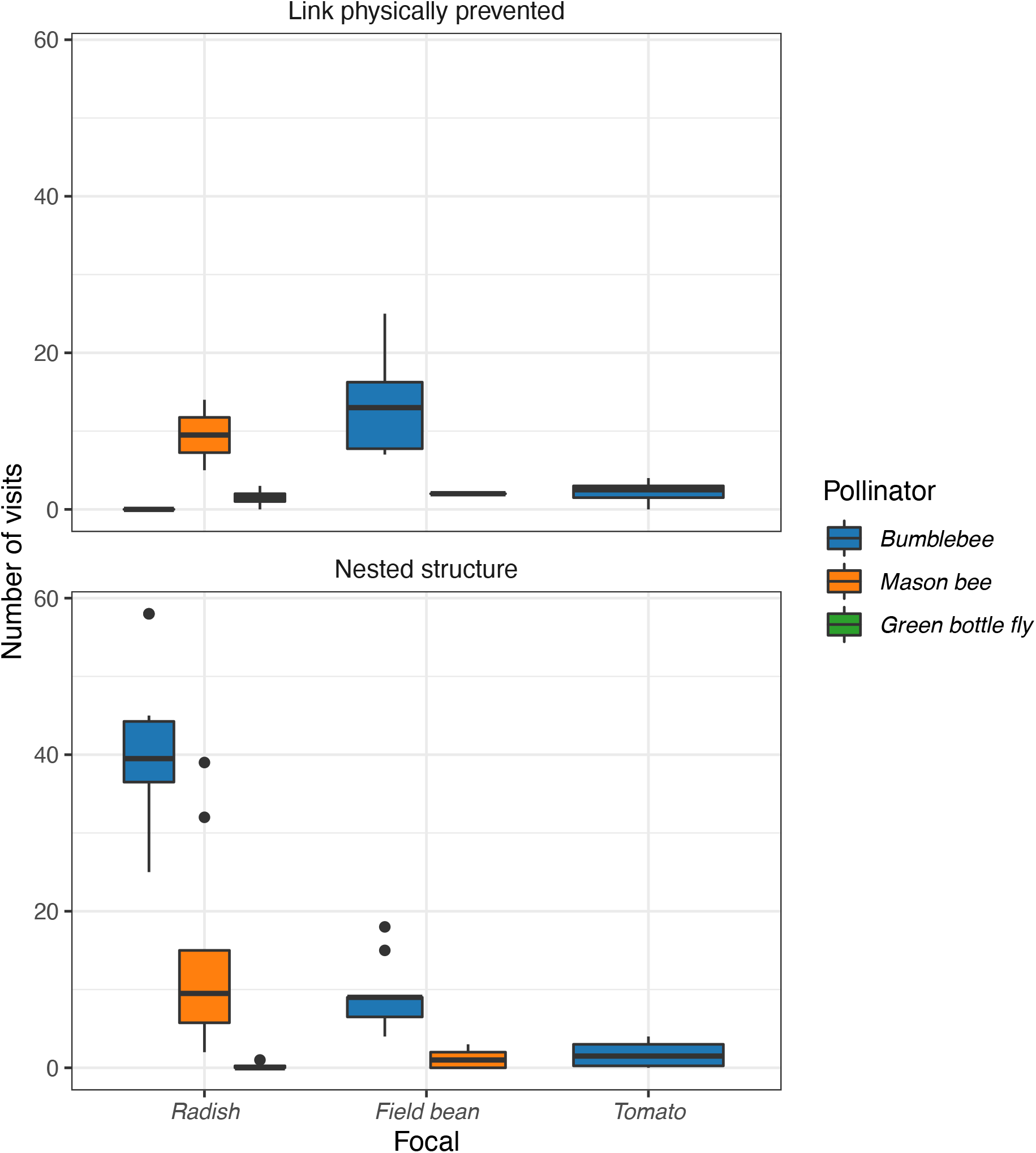
Visitation rates of each plant species by their respective pollinator(s). When the link is physically removed, the bumblebees are prevented to interact with Radishes, leading the bumblebees to interact on average more with Field beans.

### 5.2 Procrustes analysis

The interaction matrices of the non-HOIs, All HOIs, and Selected HOIs models, for both networks, were compared based on a Procrustes analysis (also called a Procrustean superimposition approach) (D. Jackson, 1995; Peres-Neto and D. A. Jackson, 2001). Briefly, a Procrustes analysis evaluates the similarity between two matrices. The procrustean analysis tests the matrices in their raw form rather than based on the corresponding distance matrices like in a mantel test (Lisboa et al., 2014). The procrustean analysis gives a correlation coefficient (Peres-Neto and D. A. Jackson, 2001) (*function: procrustes(vegan)* in R). A high correlation coefficient for the comparison of the interaction matrices A and B indicates that A and B include similar magnitude, direction, and symmetry of interactions. Here, we evaluate the similarity of two interaction matrices considering the standard variations of each interaction. Hence, the correlation coefficient between two matrices follows a distribution (see Appendix, fig.S3) according to the distributions of both matrices over the standard variations of each of their net interactions. We always compared the distribution of one interaction matrix with itself and with the distributions of the other two interaction matrices. The distribution of the correlation coefficient specific to the comparison of one matrix with itself serves as a reference to compare the other two distributions of the correlation coefficient. In ecological terms, if the distributions of correlation coefficient specific to the comparison of respectively A with A and A with B, are distinct, the matrix B includes interactions of new characteristics.

**Figure S3:**
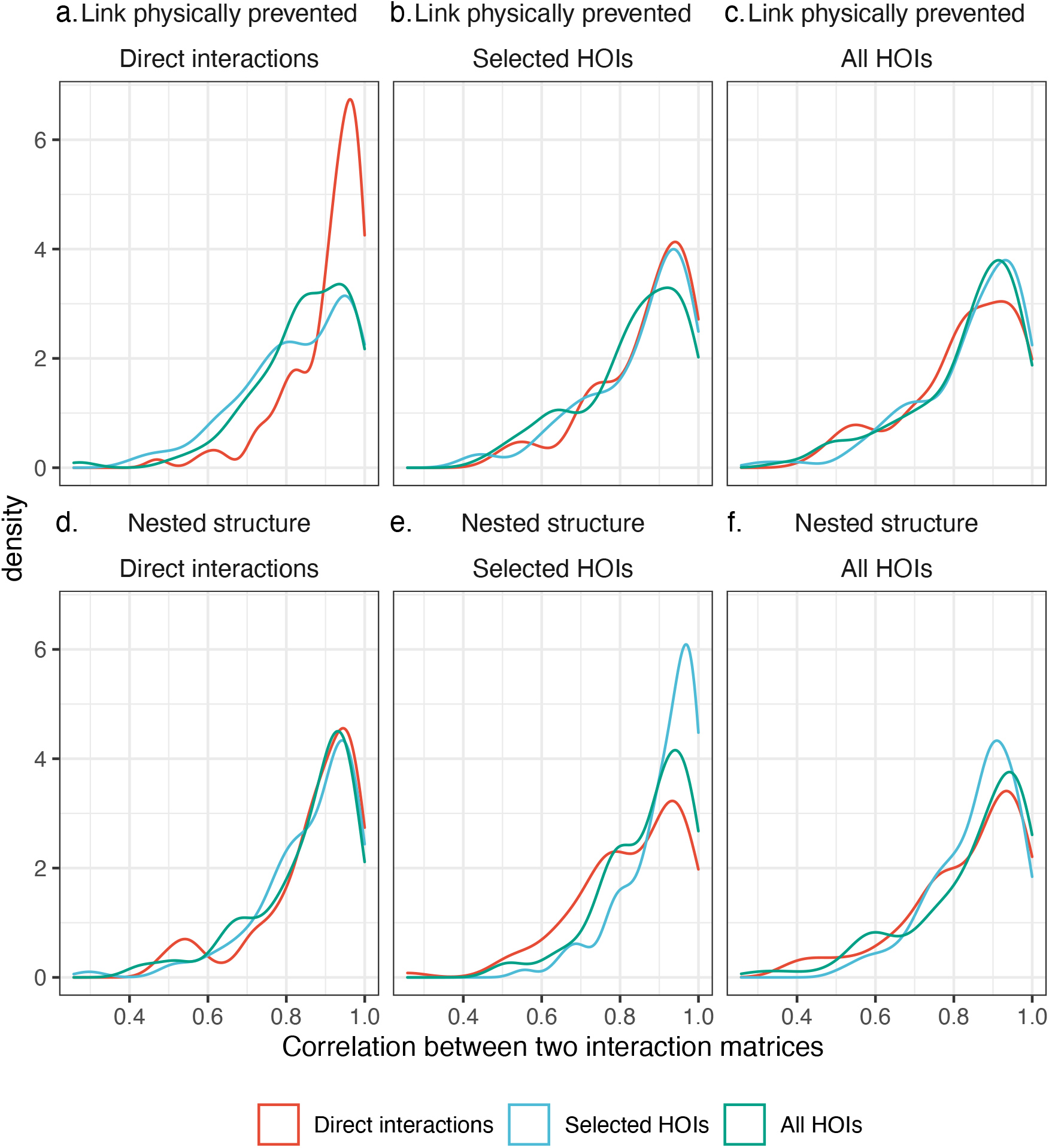
Distribution of the correlation between the distribution of a focal interaction matrix (indicated by the grid) and the distribution of another interaction matrix (indicated by the color). For instance, grid **a.** shows different correlation distributions between the comparison of the matrix with itself (here Non-HOIs or Direct interactions) and the comparison with the other model (Selected HOIs, and All HOIs).The correlation results from a Procrustes test executed over the respective distribution of each matrix according to their interaction coefficients and their standard deviations.

## 5.3 Probability Analysis

Following a structuralist probabilistic approach (Saavedra, Rohr, et al., 2017; Song, Rohr, et al., 2018), we calculate the average probability of persistence of plant species within a community A of 3 species as:

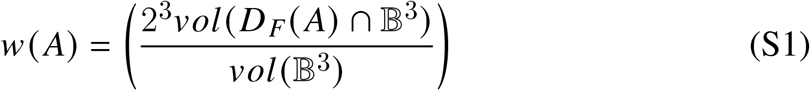

where *vol*(*BS*) is the volume of the 3-dimensional unit ball representing the parameter space of *r*, 23 normalizes the unit ball to the positive orthant, and 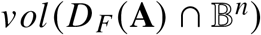 corresponds to the volume of the intersection of the feasibility domain with the unit ball. The feasibility domain is defined as 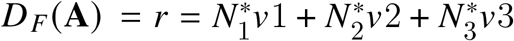, with 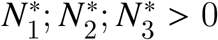, where *v_i_* is the *i*th column vector of the interaction matrix A (Song, Rohr, et al., 2018). Note that we assume that plants have positive intrinsic growth rates, i.e., *r_i_* > 0. Because we are only interested in the direction of positive r-vectors, we can consider only vectors *r* for which ‖ *r* ‖≤ 1 and normalize the size of the feasibility domain using the positive orthant of the unit ball (i.e. 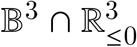). Note that we are fixing the magnitude of the r-vectors to one under the Euclidean norm (i.e. ‖ *r* ‖≤= 1), however, the analysis can be done using any norm without altering the conclusions (Rohr et al., 2016). Thus, *w*(**A**) ∈ [0; 1] is a probabilistic measure, and can be efficiently computed for even relatively large communities (Song, Rohr, et al., 2018; Song, Altermatt, et al., 2018). Ecologically, *w* (*A*) can be interpreted either as the probability of persistence of a randomly chosen species or as the expected fraction of persistent species within community A. Following the same approach (Song, Rohr, et al., 2018), we estimated the persistence probability of a given species *i* within a community (interaction matrix) A as:

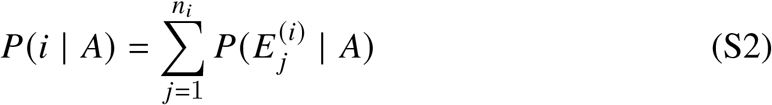

where 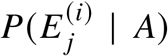 corresponds to the probability of observing the *j*-th species collection containing species *i* and *n_i_* is the total number of collections that contain species *i* starting from community A. This probability is defined as:

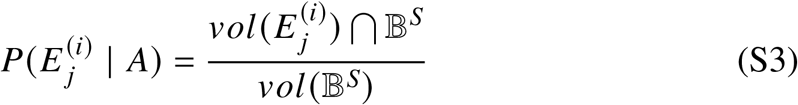

where *S* is the number of species in community A (note that in our case *S* = 3), 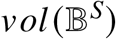 is the volume of the *S*-dimensional unit ball (i.e., the full parameter space of *r*), and 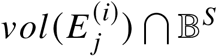 corresponds to the volume of the intersection of the domain of the collection with the unit ball. To calculate this probability we ran 100 simulations of the Lotka-Volterra dynamics using a given inferred matrix, random initial conditions, and randomly drawing r uniformly over the unit sphere ‖ *r* ‖ = 1. We assume that plants are constrained to positive intrinsic growth rates. Thus, *P*(*i* | *A*) ∈ [0; 1] and is given by the fraction of times that species *i* was found with positive abundance at equilibrium 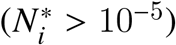.

## 5.3.1 HOIs effect on persistence probability

**Figure S4:**
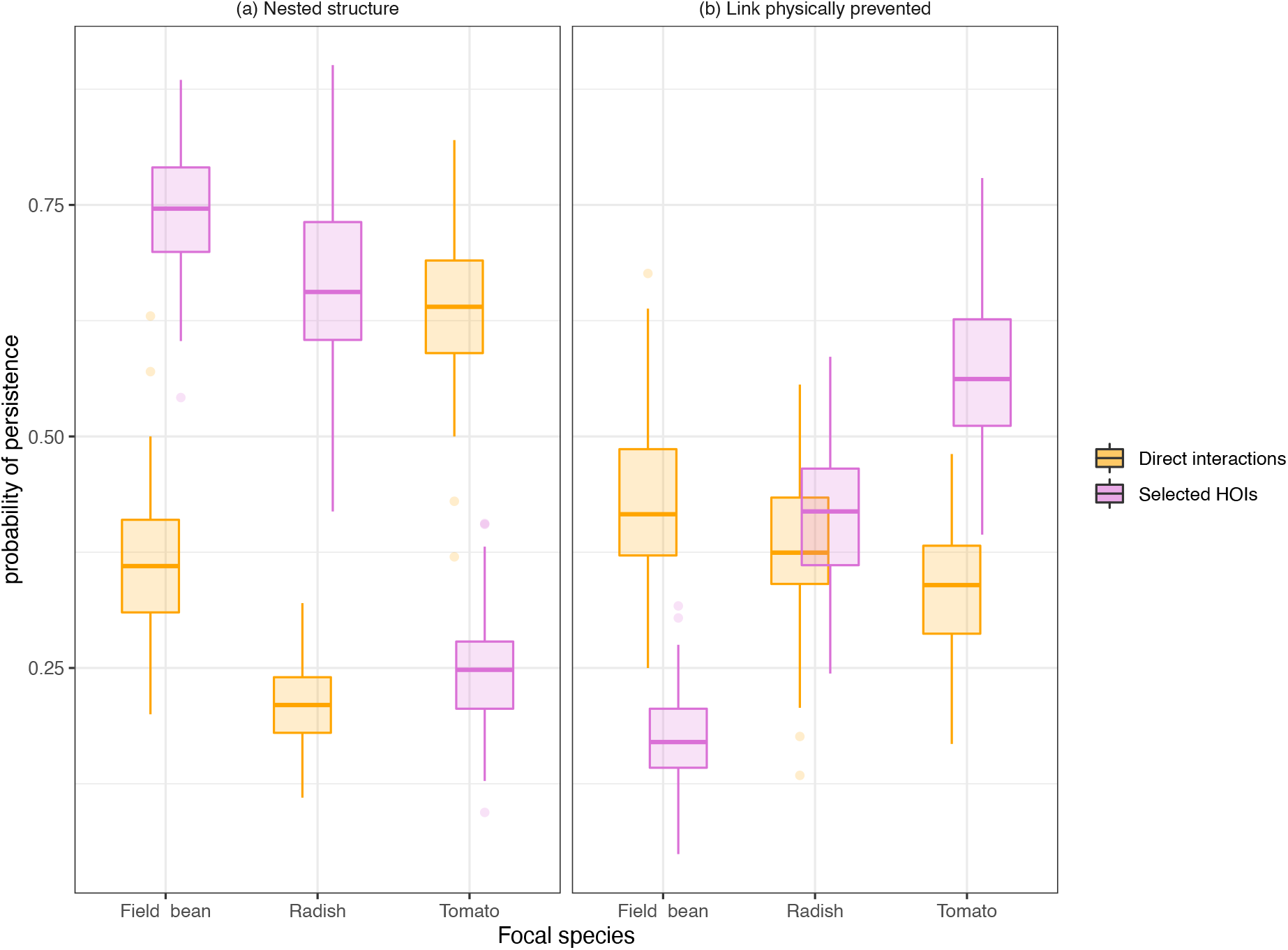
HOIs change species’ individual probability to persist. The magnitude and direction of their effect depend upon the network structure. **(b)** In a nested network, while the scenario with solely the direct interactions favor the persistence of Tomato, the HOIs scenario revert this pattern, with Tomato showing the smallest probability to persist. **(b)** When the link is physically prevented, the inclusion of HOIs drastically reduced the persistence probability of Field bean, while inducing variation in the probability of the two other plant species, but ultimately favoring Tomato. Overall, the impact of the inclusion of HOIs on species persistence probability is modulated by the network structure.

## 5.4 HOIs magnitude

**Table S1:** Effect of HOIs on the per capita sign of the interactions between plant species pairs, magnitude of HOIs compared to standard deviation of the corresponding plant direct interactions and the average number of each type of HOIs, in the Full HOIs and Selected HOIs models, under both structures.

| model | network | Positive<br>HOIs<br>plant<br>(%) | Positive<br>HOIs<br>pollinator<br>(%) | \|HOIs<br>plant\| >\|sd\|<br>of the direct<br>interaction<br>(%) | \|HOIs<br>pollinator\| >\|sd\|<br>of the direct<br>interaction<br>(%) | Number<br>of plant<br>HOIs | Number<br>of<br>pollinator<br>HOIs |
| --- | --- | --- | --- | --- | --- | --- | --- |
| All HOIs | Nested<br>structure | 0.333 | 0.444 | 0.889 | 0.333 | 3 | 3 |
| All HOIs | Link<br>physically<br>prevented | 0.333 | 0.667 | 0.778 | 0.167 | 3 | 2 |
| Selected<br>HOIs | Nested<br>structure | 0.429 | 0.5 | 1 | 0.167 | 2.33 | 2 |
| Selected<br>HOIs | Link<br>physically<br>prevented | 0.444 | 1 | 0.778 | 0.25 | 3 | 1.33 |

**Figure S5:**
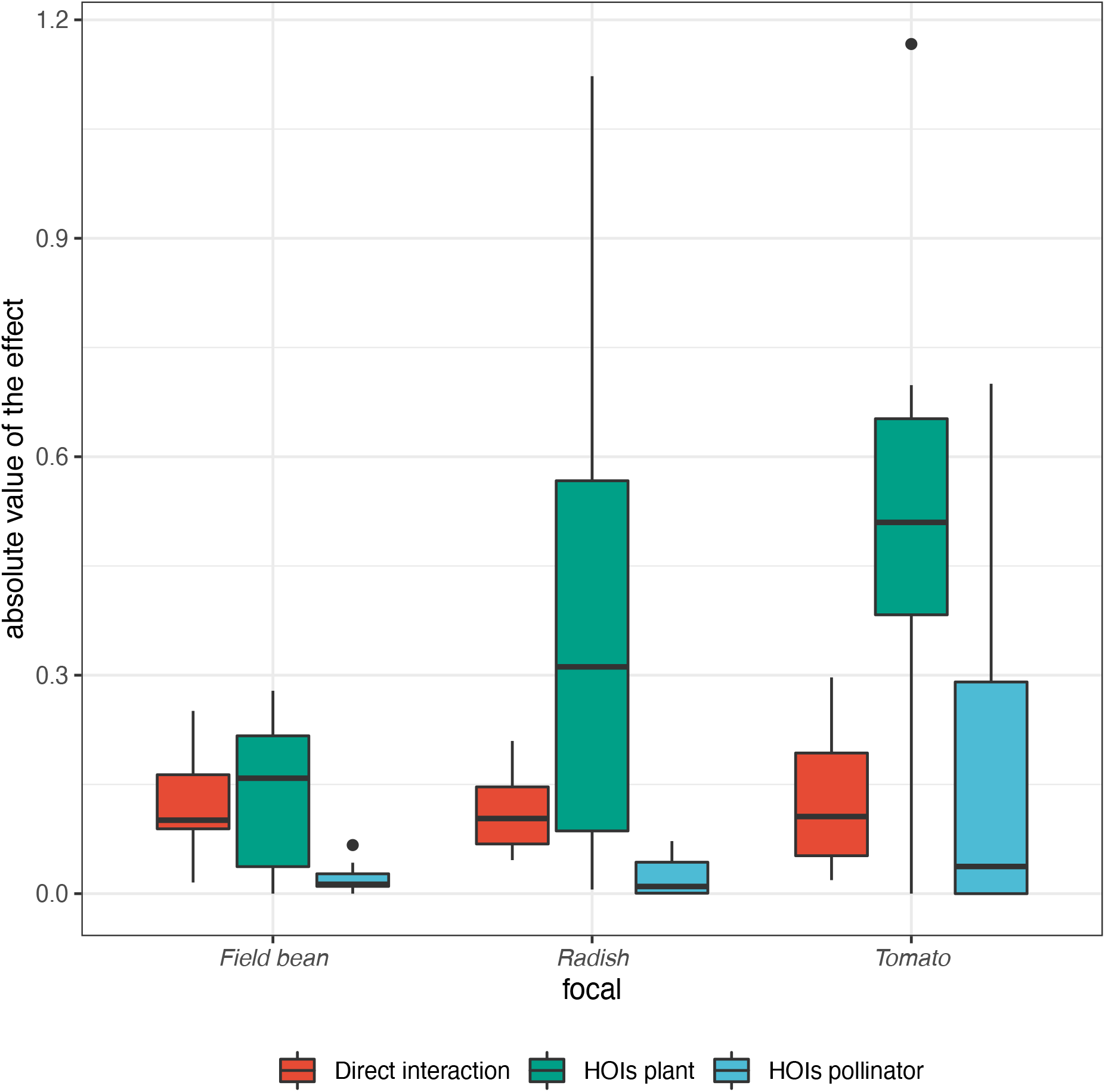
Comparison of the magnitudes of plant and pollinator HOIs with the effect of plant direct interaction (orange box-plot: *α_ij_*) for each focal species. Plant HOIs (green box-plot) are stronger than pollinator HOIs (blue box-plot). Also, the average effects of pollinator HOIs are always smaller than the average effects of plant direct interactions while they are always higher for plant HOIs. In general, the importance of HOIs appears to be species dependant.

## 5.5 Selection of models

General model:

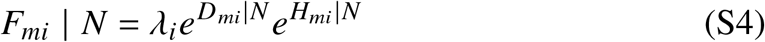

with *F_mi_* the specific fecundity of an individual *m* of the species i for a specific set *N* of neighbors, *λ_i_* the intrinsic fecundity of species *i* in the absence of competitors, and the exponential terms correspond respectively to the ensemble of pairwise interactions *D_mi_* and HOIs *H_mi_*.

Across models, the cumulative effect of the pairwise interactions is always the same:

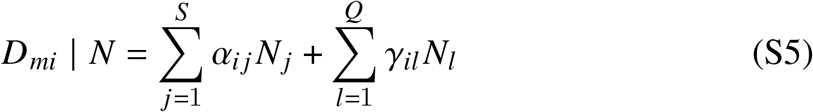

where, *N_j_* is the abundance of species j, *α_ij_* is the direct effect of species j on the focal species i, and the sum is across all the neighbor plant species *S* specific to species *i*, *γ_il_*(*i*) is the functional response of the pollinator species *l* on the focal species i, and the sum is across the pollinator species *Q* specific to species *i*. *α* can be positive or negative, capturing respectively, facilitation or competition. *γ* (*l*) can be of linear function (i.e. type I):

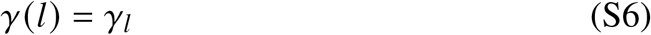

or a noninflected functions (i.e. type II):

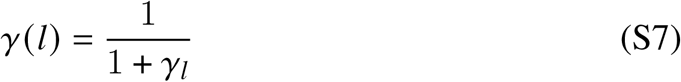

The appropriate function has been selected for each focal species by identifying the model with the smallest AIC and including at least one functional response for each pollinator of the focal species. The Table.S2 shows the functional response selected for each plant-pollinator direct interaction.

The ensemble of HOIs acting on species *i* is quantified differently according to the class of model considered.

Direct interactions:

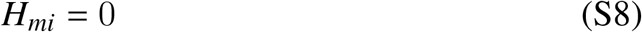

Full HOIs:

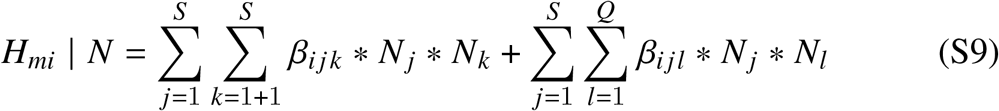

*β_ijk_* is the effect of a plant population, species *k*, on the per capita interaction between two plants, species *i* and species *j* (here *β_ijk_* is called “plant HOIs”), with *j* ≠ *k*, *β_ijl_* is the effect of a pollinator population, species *l*, on the per capita interaction between two plants, species *i* and species *j* (here *β_ijl_* is called “pollinator HOIs”). For both types of HOIs, the effect is cumulative across the plant S and/or pollinator *Q* neighbors specific to the focal species *i*. *β_ijk_* and *β_ijl_* change the nature of the resulting interaction of a competitor on the focal species *i*. For each plant species, a maximum likelihood approach allowed the estimation of each coefficient.

Similar to previous work, the decision to include the effect of HOIs on individual fitness models has been done using statistical tests that penalize for model complexity (AIC - log-likelihood) (Mayfield and Stouffer, 2017). The HOIs were selected thanks to the *dredge(MuMIn)* function in R, which used the Full HOIs model as a starting point to create a list of models ordered by statistical relevance (AIC). We filtered that list by choosing only the models with a Delta-AIC < 2 compared to the smallest AIC. Finally, we select the one model with the highest log-likelihood to minimize the AIC as *AIC* = 2*k* – 2(*log – likelihood*) with k the number of variables in the model, plus the intercept (Burnham and Anderson, 1998).

Following the philosophy of using more parsimonious models, our selected models include a lesser number of HOIs. Yet, the changes in the per capita interactions (strength and sign) and species persistence probability were not fundamentally different between the Selected model and the Full model (i.e. including all potential HOIs present on the community), and this for both network structures (Fig.S6 and Fig.S7). Two differences were noticeable. First, the inclusion of all potential HOIs increases the standard variation respective to the nine per capita interactions (Fig.S6). Second, contrary to a full model, the selected model includes a positive probability of persistence of the three plant species community in a nested structure (Fig.S7). Also, at a community level approach, the Procrustes analysis did show distributions of correlation between species interactions matrices following similar trends for the Selected and Full HOIs models (appendix 5.2, Fig.S3). Thus, despite the two previously mentioned differences, both models point out similar ecological conclusions, in favor of the inclusion of relevant multitrophic HOIs.

---

**Figure S6:**
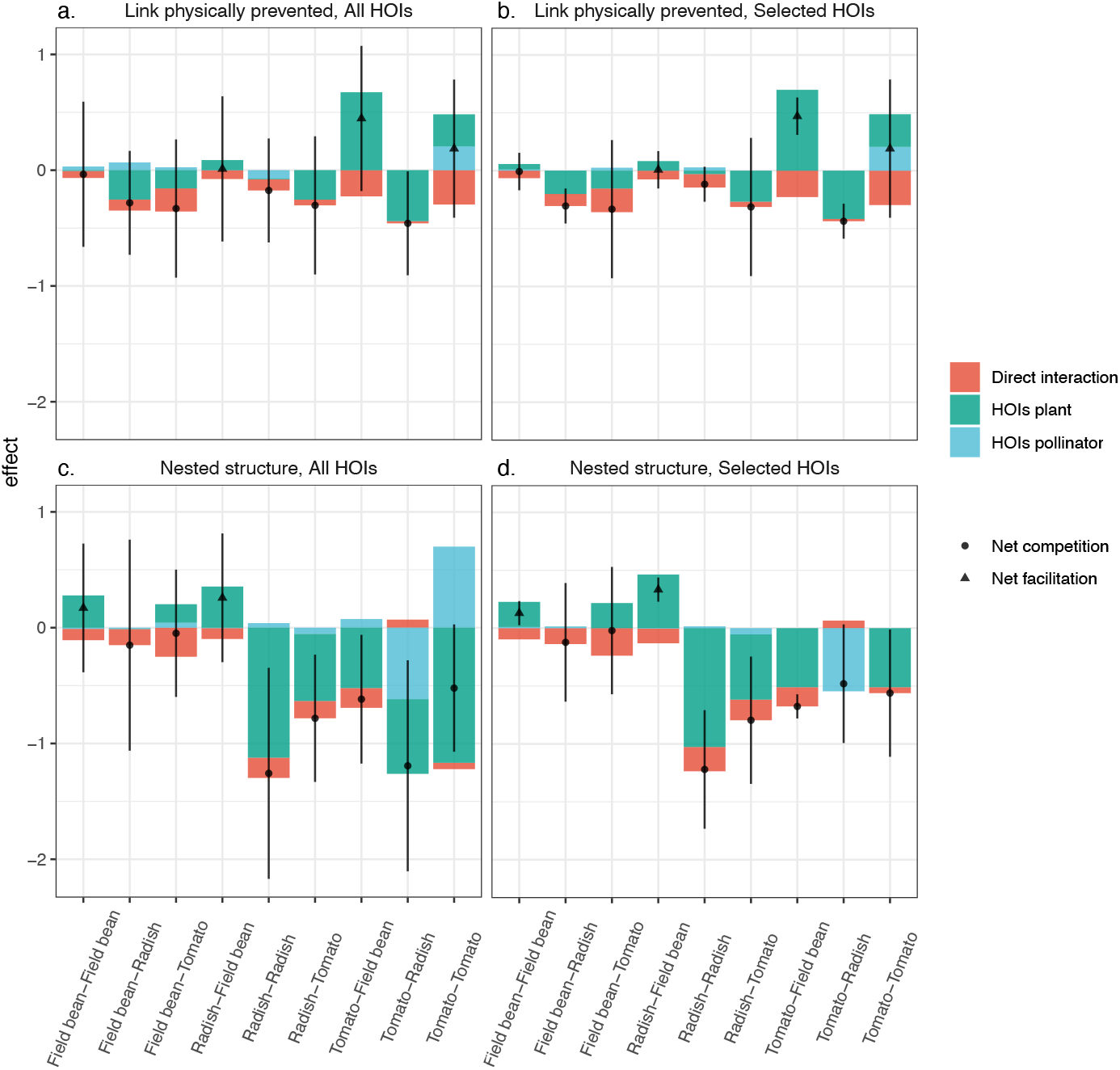
The inclusion of only the statistically selected HOIs reduces the standard variation of the realized per capita interaction. Moreover, in a nested structure, the magnitude of HOIs pollinators increases when the model is forced to estimate them. The nine direct plant interactions for the four scenarios are divided in terms of direct interaction(s) (red), HOIs plant (blue), and HOIs pollinator (green). The dots represent the sum of all interactions (i.e. the realized per capita interaction), that is the value of the strength between the two corresponding plant species. The dot is circular if the value is negative (i.e. net competition) and triangular if the value is positive (i.e. net facilitation). Each dot is associated with its standard deviation.

**Figure S7:**
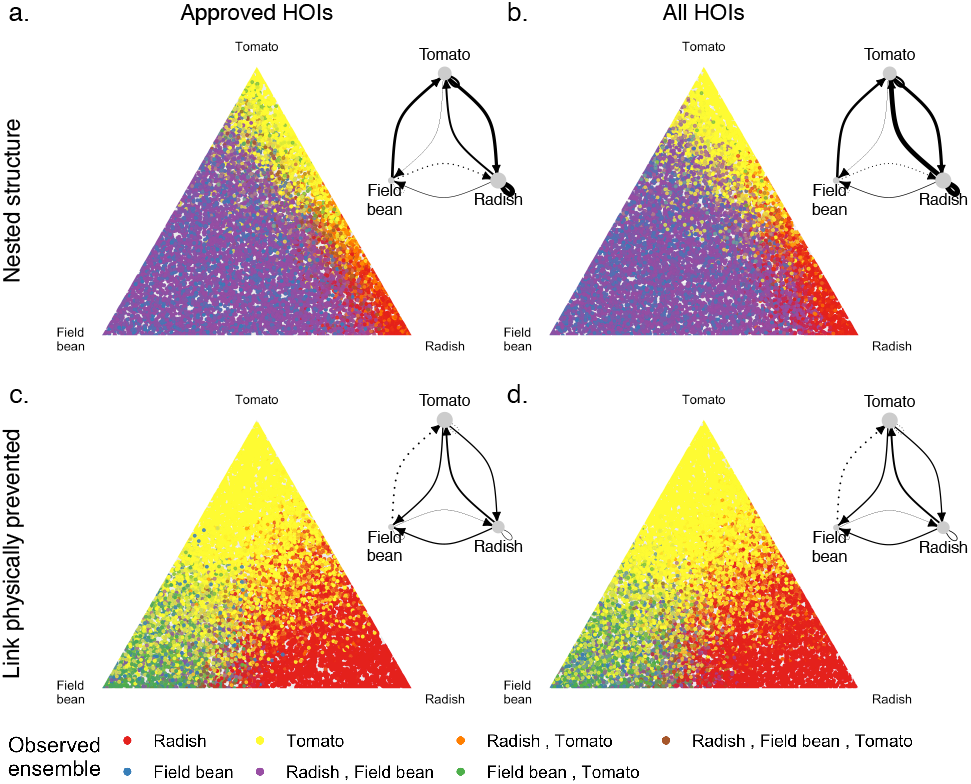
The probability of species persistence does not change between a model with all HOIs or only the statistically selected HOIs. A noticeable difference is the inclusion of the three-species combination in the selected HOIs scenario, under a nested structure. The percentage of persistence probability of each plant species individually, and in all pairs and triplet combinations is proportional to the fill of the triangle under the corresponding color. Each triangle is associated with its corresponding diagram of species interactions (solid lines = net competitive interaction, dotted lines = net facilitation and the width of the line is proportional to the absolute value of the strength of the interaction).

**Table S2:** Coefficients of the direct interactions and the HOIs included in the selected model for each plant focal and network structure. The total number of parameters is given at the beginning of the row. Note that a negative coefficient of a type I pollinator functional response indicates a hump-shaped function with a positive effect of visitation on seed set at low visitation rates values, and a slightly negative effect at very high values

| **species** | **model** | **network** | **df** | **AIC** | **logLik** | **Intercept** | **year2017** |
| --- | --- | --- | --- | --- | --- | --- | --- |
| Field bean | with HOIs | no link | 12 | 904.011 | -440.005 | 1.080 | 0.127 |
| Field bean | no HOIs | no link | 8 | 908.493 | -446.247 | 1.115 | 0.147 |
| Radish | with HOIs | no link | 12 | 2617.313 | -1296.656 | 2.810 | 0.853 |
| Radish | no HOIs | no link | 8 | 2617.111 | -1300.556 | 2.923 | 0.896 |
| Tomato | with HOIs | no link | 10 | 807.528 | -393.764 | 4.448 | 1.131 |
| Tomato | no HOIs | no link | 7 | 809.142 | -397.571 | 4.414 | 0.879 |
| Field bean | with HOIs | with link | 11 | 1281.968 | -629.984 | 1.170 | 0.147 |
| Field bean | no HOIs | with link | 8 | 1284.510 | -634.255 | 1.233 | 0.174 |
| Radish | with HOIs | with link | 15 | 2601.208 | -1285.604 | 1.750 | 1.147 |
| Radish | no HOIs | with link | 9 | 2601.747 | -1291.874 | 1.757 | 1.166 |
| Tomato | with HOIs | with link | 9 | 635.998 | -308.999 | 4.285 | 0.147 |
| Tomato | no HOIs | with link | 7 | 635.213 | -310.606 | 4.319 | 0.057 |

| **species** | **model** | **network** | **df** | **Radish** | **Field bean** | **Tomato** |
| --- | --- | --- | --- | --- | --- | --- |
| Field bean | with HOIs | no link | 12 | -0.104 | -0.065 | -0.204 |
| Field bean | no HOIs | no link | 8 | -0.115 | 0.015 | -0.094 |
| Radish | with HOIs | no link | 12 | -0.116 | -0.075 | -0.046 |
| Radish | no HOIs | no link | 8 | -0.048 | -0.066 | -0.109 |
| Tomato | with HOIs | no link | 10 | -0.020 | -0.228 | -0.297 |
| Tomato | no HOIs | no link | 7 | -0.030 | -0.198 | -0.140 |
| Field bean | with HOIs | with link | 11 | -0.138 | -0.097 | -0.238 |
| Field bean | no HOIs | with link | 8 | -0.087 | -0.051 | -0.172 |
| Radish | with HOIs | with link | 15 | -0.210 | -0.127 | -0.178 |
| Radish | no HOIs | with link | 9 | -0.066 | -0.138 | -0.160 |
| Tomato | with HOIs | with link | 9 | 0.065 | -0.166 | -0.051 |
| Tomato | no HOIs | with link | 7 | 0.042 | -0.179 | -0.072 |

| **species** | **model** | **network** | **df** | **Bumblebee** | **Fly** | **Bee** | **I(1/(1 + Bumblebee))** | **I(1/(1 + Fly))** | **I(1/(1 + Bee))** |
| --- | --- | --- | --- | --- | --- | --- | --- | --- | --- |
| Field bean | with HOIs | no link | 12 | NA | NA | NA | 0.325 | NA | 0.382 |
| Field bean | no HOIs | no link | 8 | NA | NA | NA | 0.060 | NA | 0.512 |
| Radish | with HOIs | no link | 12 | NA | -0.078 | NA | NA | NA | 0.407 |
| Radish | no HOIs | no link | 8 | NA | -0.091 | NA | NA | NA | 0.260 |
| Tomato | with HOIs | no link | 10 | -0.537 | NA | NA | NA | NA | NA |
| Tomato | no HOIs | no link | 7 | -0.300 | NA | NA | NA | NA | NA |
| Field bean | with HOIs | with link | 11 | NA | NA | -0.165 | 0.672 | NA | NA |
| Field bean | no HOIs | with link | 8 | NA | NA | -0.152 | 0.522 | NA | NA |
| Radish | with HOIs | with link | 15 | -0.014 | NA | NA | NA | 0.947 | 0.674 |
| Radish | no HOIs | with link | 9 | -0.010 | NA | NA | NA | 0.892 | 0.643 |
| Tomato | with HOIs | with link | 9 | -0.568 | NA | NA | NA | NA | NA |
| Tomato | no HOIs | with link | 7 | -0.628 | NA | NA | NA | NA | NA |

| **species** | **model** | **network** | **df** | **Radish:Tomato** | **Tomato:Field bean** | **Radish:Field bean** |
| --- | --- | --- | --- | --- | --- | --- |
| Field bean | with HOIs | no link | 12 | -0.201 | 0.047 | NA |
| Field bean | no HOIs | no link | 8 | NA | NA | NA |
| Radish | with HOIs | no link | 12 | -0.190 | -0.078 | 0.161 |
| Radish | no HOIs | no link | 8 | NA | NA | NA |
| Tomato | with HOIs | no link | 10 | -0.417 | 0.698 | NA |
| Tomato | no HOIs | no link | 7 | NA | NA | NA |
| Field bean | with HOIs | with link | 11 | NA | 0.217 | NA |
| Field bean | no HOIs | with link | 8 | NA | NA | NA |
| Radish | with HOIs | with link | 15 | -1.026 | 0.463 | NA |
| Radish | no HOIs | with link | 9 | NA | NA | NA |
| Tomato | with HOIs | with link | 9 | NA | -0.510 | NA |
| Tomato | no HOIs | with link | 7 | NA | NA | NA |

| **species** | **model** | **network** | **df** | **Bumblebee:Radish** | **Bumblebee:Tomato** | **Bumblebee:Field bean** |
| --- | --- | --- | --- | --- | --- | --- |
| Field bean | with HOIs | no link | 12 | NA | 0.026 | 0.010 |
| Field bean | no HOIs | no link | 8 | NA | NA | NA |
| Radish | with HOIs | no link | 12 | NA | NA | NA |
| Radish | no HOIs | no link | 8 | NA | NA | NA |
| Tomato | with HOIs | no link | 10 | NA | 0.206 | NA |
| Tomato | no HOIs | no link | 7 | NA | NA | NA |
| Field bean | with HOIs | with link | 11 | 0.015 | NA | 0.010 |
| Field bean | no HOIs | with link | 8 | NA | NA | NA |
| Radish | with HOIs | with link | 15 | 0.007 | -0.054 | NA |
| Radish | no HOIs | with link | 9 | NA | NA | NA |
| Tomato | with HOIs | with link | 9 | -0.545 | NA | NA |
| Tomato | no HOIs | with link | 7 | NA | NA | NA |

| **species** | **model** | **network** | **df** | **Fly:Radish** | **Bee:Field bean** | **Bee:Radish** |
| --- | --- | --- | --- | --- | --- | --- |
| Field bean | with HOIs | no link | 12 | NA | NA | NA |
| Field bean | no HOIs | no link | 8 | NA | NA | NA |
| Radish | with HOIs | no link | 12 | NA | NA | 0.028 |
| Radish | no HOIs | no link | 8 | NA | NA | NA |
| Tomato | with HOIs | no link | 10 | NA | NA | NA |
| Tomato | no HOIs | no link | 7 | NA | NA | NA |
| Field bean | with HOIs | with link | 11 | NA | NA | NA |
| Field bean | no HOIs | with link | 8 | NA | NA | NA |
| Radish | with HOIs | with link | 15 | NA | -0.004 | 0.009 |
| Radish | no HOIs | with link | 9 | NA | NA | NA |
| Tomato | with HOIs | with link | 9 | NA | NA | NA |
| Tomato | no HOIs | with link | 7 | NA | NA | NA |

## References

Adler, P. B., HilleRisLambers, J., and Levine, J. M. (2007). A niche for neutrality. In: Ecology Letters 10.2, pp. 95–104.

Allesina, S. and Levine, J. M. (2011). A competitive network theory of species diversity. In: Proceedings of the National Academy of Sciences of the United States of America 108.14, pp. 5638–5642.

Bairey, E., Kelsic, E. D., and Kishony, R. (2016). High-order species interactions shape ecosystem diversity. In: Nature Communications 7, pp. 1–7.

Barabás, G., J. Michalska-Smith, M., and Allesina, S. (2016). The effect of intra-and interspecific competition on coexistence in multispecies communities. In: The American Naturalist 188.1, E1–E12.

Bartley, T. J. et al. (2019). Food web rewiring in a changing world. In: Nature ecology & evolution 3.3, pp. 345–354.

Bartomeus, I., Saavedra, S., Rohr, R. P., and Godoy, O. (2021). Experimental evidence of the importance of multitrophic structure for species persistence. In: Proceedings of the National Academy of Sciences 118.12.

Bastolla, U., Fortuna, M. A., Pascual-García, A., Ferrera, A., Luque, B., and Bascompte, J. (2009). The architecture of mutualistic networks minimizes competition and increases biodiversity. In: Nature 458.7241, pp. 1018–1020.

Beckerman, A. P., Uriarte, M., and Schmitz, O. J. (1997). Experimental evidence for a behavior-mediated trophic cascade in a terrestrial food chain. In: Proceedings of the National Academy of Sciences of the United States of America 94.20, pp. 10735–10738.

Billick, I. and Case, T. J. (1994). Higher Order Interactions in Ecological Communities: What Are They and How Can They be Detected? In: Ecology 75.6, pp. 1530–1543.

Burkle, L. A. and Runyon, J. B. (2019). Floral volatiles structure plant–pollinator interactions in a diverse community across the growing season. In: Functional Ecology 33.11. Ed. by J. Manson, pp. 2116–2129.

Burnham, K. P. and Anderson, D. R. (1998). Model Selection and Multimodel Inference: A Practical Information-Theoretic Approach. Springer, New York, NY.

CaraDonna, P. J. et al. (2017). Interaction rewiring and the rapid turnover of plant-pollinator networks. In: Ecology letters 20.3, pp. 385–394.

Case, T. J. and Bender, E. A. (1981). Testing for Higher Order Interactions. In: The American Naturalist 118.6, pp. 920–929.

Cattin, M.-F., Bersier, L.-F., Banašek-Richter, C., Baltensperger, R., and Gabriel, J.-P. (2004). Phylogenetic constraints and adaptation explain food-web structure. In: Nature 427.6977, pp. 835–839.

Chesson, P. (2000). General Theory of Competitive Coexistence in Spatially - Varying Environments. In: Theoretical Population Biology 58.3, pp. 211–237.

Godoy, O., Bartomeus, I., Rohr, R. P., and Saavedra, S. (2018). Towards the Integration of Niche and Network Theories. In: Trends in Ecology & Evolution 33.4, pp. 287–300.

Godoy, O. and Levine, J. M. (Mar. 2014). Phenology effects on invasion success: insights from coupling field experiments to coexistence theory. In: Ecology 95.3, pp. 726–736.

Godoy, O., Stouffer, D. B., Kraft, N. J. B., and Levine, J. M. (2017). Intransitivity is infrequent and fails to promote annual plant coexistence without pairwise niche differences. In: Ecology 98.5, pp. 1193–1200.

Grilli, J., Barabás, G., Michalska-Smith, M. J., and Allesina, S. (2017). Higher-order interactions stabilize dynamics in competitive network models. In: Nature 548.7666, pp. 210–213.

Jackson, D. (1995). PROTEST: A PROcrustean Randomization TEST of community environment concordance. Vol. 2. 3, pp. 297–303.

Johnson, C. A. (2021). How mutualisms influence the coexistence of competing species. Wiley Online Library, e03346.

Kleinhesselink, A. R., Kraft, N. J., and Levine, J. M. (2019). Mechanisms underlying higher order interactions: from quantitative definitions to ecological processes. Tech. rep.

Lanuza, J. B., Bartomeus, I., and Godoy, O. (2018). Opposing effects of floral visitors and soil conditions on the determinants of competitive outcomes maintain species diversity in heterogeneous landscapes. In: Ecology Letters 21.6. Ed. by J. M. Gómez, pp. 865–874.

Letten, A. D. and Stouffer, D. B. (2019). The mechanistic basis for higher-order interactions and non-additivity in competitive communities. In: Ecology Letters 22.3, pp. 423–436.

Levine, J. M., Bascompte, J., Adler, P. B., and Allesina, S. (2017). Beyond pairwise mechanisms of species coexistence in complex communities. In: Nature 546.7656, pp. 56–64.

Li, Y., Bearup, D., and Liao, J. (2020). Habitat loss alters effects of intransitive higher-order competition on biodiversity: a new metapopulation framework. In: Proceedings of the Royal Society B.

Li, Y., Mayfield, M. M., et al. (2021). Beyond direct neighbourhood effects: higher-order interactions improve modelling and predicting tree survival and growth. In: National Science Review 8.5, pp. 169–178.

Lisboa, F. J. G. et al. (2014). Much beyond Mantel: Bringing procrustes association metric to the plant and soil ecologist’s toolbox. In: PLoS ONE 9.6, pp. 1–9.

Martyn, T. E., Stouffer, D. B., Godoy, O., Bartomeus, I., Pastore, A. I., and Mayfield, M. M. (2021). Identifying âUsefulâ Fitness Models: Balancing the Benefits of Added Complexity with Realistic Data Requirements in Models of Individual Plant Fitness. In: The American Naturalist 197.4. PMID: 33755538, pp. 415–433.

May, R. M. and Leonard, W. J. (1975). Nonlinear Aspects of Competition Between Three Species. In: SIAM Journal on Applied Mathematics 29.2, pp. 243–253.

Mayfield, M. M. and Stouffer, D. B. (2017). Higher-order interactions capture unexplained complexity in diverse communities. In: Nature Ecology & Evolution 1.3, p. 62.

McCann, K., Hastings, A., and Huxel, G. R. (1998). Weak trophic interactions and the balance of nature. In: Nature 395.6704, pp. 794–798.

Montoya, J. M., Pimm, S. L., and Solé, R. V. (2006). Ecological networks and their fragility. Vol. 442.7100. Nature Publishing Group, pp. 259–264.

Morales-Castilla, I., Matias, M. G., Gravel, D., and Araújo, M. B. (2015). Inferring biotic interactions from proxies. In: Trends in ecology & evolution 30.6, pp. 347–356.

Morin, P. J., Lawler, S. P., and Johnson, E. A. (1988). Competition Between Aquatic Insects and Vertebrates: Interaction Strength and Higher Order. Tech. rep. 5, pp. 1401–1409.

Morris, W. F., Vazquez, D. P., and Chacoff, N. P. (2010). Benefit and cost curves for typical pollination mutualisms. In: Ecology 91.5, pp. 1276–1285.

Neutel, A.-M. et al. (2007). Reconciling complexity with stability in naturally assembling food webs. In: Nature 449.7162, pp. 599–602.

Paterson, R. A., Dick, J. T. A., Pritchard, D. W., Ennis, M., Hatcher, M. J., and Dunn, A. M. (2015). Predicting invasive species impacts: a community module functional response approach reveals context dependencies. In: Journal of Animal Ecology 84.2. Ed. by B. Woodcock, pp. 453–463.

Peres-Neto, P. R. and Jackson, D. A. (2001). How well do multivariate data sets match? The advantages of a procrustean superimposition approach over the Mantel test. In: Oecologia 129.2, pp. 169–178.

Perez-Ramos, I. M., Matias, L., Gomez-Aparicio, L., and Godoy, O. (2019). Functional traits and phenotypic plasticity modulate species coexistence across contrasting climatic conditions. In: Nature Communications 10.1, p. 2555.

Rohr, R. P. et al. (2016). Persist or produce: A community trade-off tuned by species evenness. In: American Naturalist 188.4, pp. 411–422.

Rumeu, B., Álvarez-Villanueva, M., Arroyo, J. M., and González-Varo, J. P. (2019). Interspecific competition for frugivores: population-level seed dispersal in contrasting fruiting communities. In: Oecologia 190.3, pp. 605–617.

Saavedra, S., Medeiros, L. P., and AlAdwani, M. (2020). Structural forecasting of species persistence under changing environments. In: Ecology Letters 23.

Saavedra, S., Rohr, R. P., Bascompte, J., Godoy, O., Kraft, N. J. B., and Levine, J. M. (2017). A structural approach for understanding multispecies coexistence. In: Ecological Monographs 87.3, pp. 470–486.

Singh, P. and Baruah, G. (2019). Higher order interactions and coexistence theory. In: bioRxiv.

Song, C., Altermatt, F., Pearse, I., and Saavedra, S. (2018). Structural changes within trophic levels are constrained by within-family assembly rules at lower trophic levels. In: Ecology Letters 21.8, pp. 1221–1228.

Song, C., Rohr, R. P., and Saavedra, S. (2018). A guideline to study the feasibility domain of multi-trophic and changing ecological communities. In: Journal of Theoretical Biology 450, pp. 30–36.

Verdú, M. and Valiente-Banuet, A. (2008). The nested assembly of plant facilitation networks prevents species extinctions. In: The American Naturalist 172.6, pp. 751–760.

Vivaldo, G., Masi, E., Taiti, C., Caldarelli, G., and Mancuso, S. (2017). The network of plants volatile organic compounds. In: Scientific Reports 7.1, pp. 1–18.

Winfree, R., Williams, N. M., Gaines, H., Ascher, J. S., and Kremen, C. (2008). Wild bee pollinators provide the majority of crop visitation across land-use gradients in New Jersey and Pennsylvania, USA. In: Journal of applied ecology 45.3, pp. 793–802.

Wissinger, S. and Jill, M. (1993). Intraguild predation and competition between larval dragonflies: direct and indirect effects on shared prey. In: Ecological society of America 74.1, pp. 207–218.

Worthen, W. B. and Moore, J. L. (1991). Higher-order interactions and indirect effects: a resolution using laboratory Drosophila communities. In: The American Naturalist 138.5, pp. 1092–1104.

## References

D. Jackson (1995). PROTEST: A PROcrustean Randomization TEST of community environment concordance. Vol. 2. 3, pp. 297–303

F. J. G. Lisboa et al. (2014). Much beyond Mantel: Bringing procrustes association metric to the plant and soil ecologist’s toolbox. In: PLoS ONE 9.6, pp. 1–9

P. R. Peres-Neto and D. A. Jackson (2001). How well do multivariate data sets match? The advantages of a procrustean superimposition approach over the Mantel test. In: Oecologia 129.2, pp. 169–178

## References

R. P. Rohr et al. (2016). Persist or produce: A community trade-off tuned by species evenness. In: American Naturalist 188.4, pp. 411–422

C. Song, R. P. Rohr, et al. (2018). A guideline to study the feasibility domain of multi-trophic and changing ecological communities. In: Journal of Theoretical Biology 450, pp. 30–36

S. Saavedra, R. P. Rohr, et al. (2017). A structural approach for understanding multispecies coexistence. In: Ecological Monographs 87.3, pp. 470–486

C. Song, F. Altermatt, et al. (2018). Structural changes within trophic levels are constrained by within-family assembly rules at lower trophic levels. In: Ecology Letters 21.8, pp. 1221–1228

## References

K. P. Burnham and D. R. Anderson (1998). Model Selection and Multimodel Inference: A Practical Information-Theoretic Approach. Springer, New York, NY.

M. M. Mayfield and D. B. Stouffer (2017). Higher-order interactions capture unexplained complexity in diverse communities. In: Nature Ecology & Evolution 1.3, p. 62

